# A glyphosate-based herbicide selects for genetic changes while retaining within-species diversity in a freshwater bacterioplankton community

**DOI:** 10.1101/2024.09.17.613573

**Authors:** Emma Derrick, Naíla Barbosa da Costa, Rowan D. H. Barrett, B. Jesse Shapiro

## Abstract

Bacterial populations evolve rapidly in the lab when faced with experimentally-applied selective pressures. Yet how bacteria evolve in nature, in more complex multi-species communities, is both challenging to study and essential to our understanding of ecosystem responses to rapid anthropogenic change. It has been theorized that selection purges within-species diversity in genome-wide selective sweeps, but the prevalence of such sweeps in response to known selective pressures in nature remains unclear. To track bacterial evolution in a semi-natural context, we applied Roundup, a glyphosate-based herbicide (GBH) as a selective pressure to 1000 L ponds containing bacterioplankton communities from a pristine lake. Using metagenomic analyses, we found that GBH treatment substantially affected community diversity, reducing species richness twofold, but did not consistently purge within-species genetic diversity over the four weeks of the experiment. We identified several functional categories of genes targeted by GBH selection across 11 different species of bacteria. There was no evidence for selection on the enzyme targeted by glyphosate, which interferes with amino acid synthesis; however genes involved more broadly in amino acid transport and metabolism were more likely to experience changes in allele frequency, particularly in inferred GBH-sensitive species. Together, these results show how environmental change can rapidly affect bacterial community structure while leaving within-species diversity largely intact. Even without evident genome-wide selective sweeps, we identify consistent genetic targets of selection, pointing to alternative mechanisms of GBH resistance in nature, and suggesting a role for soft or gene-specific selective sweeps in adaptation.

## Introduction

Microbial communities and the populations within them form the foundation of all ecosystems on Earth (1). Many studies have focused on how natural microbial communities change on an ecological level, with species changing in abundance in response to perturbations from environmental stressors (2–6). It is becoming increasingly evident that microbial populations within communities can also evolve on the same time scales as these ecological changes (7–9). Rapid evolutionary change, for example of pathogens within infected patients, can have consequences for virulence and disease persistence (10–13). More generally, within-species diversity is important for the functioning and stability of bacterial communities (14–16). For example, evolutionary diversification of a focal species can impact relative abundances of other species within the community (17). Conversely, higher community diversity may promote or constrain the diversification of species within a community (18). In the human gut, community diversity promotes within-species diversity over time scales of a few months (“diversity begets diversity”), until niches are filled (8). In other more diverse communities such as freshwater, sediments, and soil microbiomes, the “diversity begets diversity” effect is negligible, presumably because most niches are already filled (19). Similarly, natural compost communities challenged with copper stress showed independent evolutionary and ecological changes, with no detectable interaction between the two (7). In all cases, ecological and evolutionary changes within microbial communities, along with the interactions in some cases, are expected to affect ecosystems functions.

What types of evolutionary changes can occur within a community? Under the stable ecotype model, selection on adaptive mutations can result in a genome-wide selective sweep, in which a genome with an adaptive allele expands clonally (with relatively little recombination) and purges genetic diversity from the population (20–22). Alternatively, if recombination is high, an adaptive allele can be exchanged by horizontal gene transfer and spread through the population in a gene-specific sweep, purging diversity in a region of the genome while maintaining genome-wide diversity (23, 24). Or, if an adaptive allele is originally present in the population on multiple genetic backgrounds, then, during a sweep, multiple strains carrying the adaptive allele may increase in frequency, resulting in a soft sweep (25). Both soft and gene-specific selective sweeps allow specific genes targeted by selection to adapt without purging diversity genome-wide. They differ in that soft sweeps occur when recurrent adaptive mutations occur on different genomic backgrounds (lineages or strains) while gene-specific sweeps require high rates of recombination relative to selection.

In nature, there has been little evidence for pervasive genome-wide selective sweeps. Even in simplified laboratory evolution experiments where genome-wide selective sweeps are theorized to be more likely, diversity is often maintained in the population by diverse genetic targets of selection and clonal interference (26, 27). In natural environments such as the human gut, high rates of recombination allow for the exchange of genes within and between species (28, 29), potentially promoting gene-specific rather than genome-wide sweeps (30). Studies of natural populations of bacteria in freshwater lakes have identified evolutionary patterns consistent with sweeps, but it remains unclear if these were driven by selection or genetic drift. Bendall et al. (31) observed a gradual loss in diversity over nine years in one population of bacteria, consistent with a genome-wide sweep, as well as other populations with low diversity in small genomic regions, consistent with past gene-specific sweeps. More recently, Rower et al. (32) quantified changes in species abundance and diversity throughout a 20-year time series and identified an association with seasonal changes, as well as one possible soft sweep in *Nanopelagicus*. While these studies suggest that selective sweeps are occurring in natural bacterioplankton populations, neither could attribute sweeps to a known selective pressure.

Agrochemical pollution is an important selective pressure relevant to soil and aquatic microbial communities. There is growing concern over agrochemicals entering freshwater from runoff and leaching from agricultural land (33). Glyphosate-based herbicides (GBHs) are the most commonly used herbicides worldwide, and while the active ingredient glyphosate is thought to strongly bind to soil (34), it may also run off into rivers, streams, and lakes. Regulations in Canada limit the concentration of glyphosate permitted for chronic (< 800 µg/L) and acute (< 27000 µg/L) aquatic contamination (34). However, these guidelines are based on toxicity to eukaryotes, and ignore potential effects on bacteria. Previous studies have shown GBH alters the composition of bacterial communities in water (2) and honeybees (5). Further, GBH can cross-select for antibiotic resistance genes in soil (35) and aquatic (36) communities.

GBHs inhibit plant growth by interfering with the shikimate pathway and preventing downstream synthesis of essential aromatic amino acids (37). Glyphosate prevents the conversion of shikimate-3-phosphate (S3P) and phosphoenolpyruvate (PEP) into 5-enolpyruvylshikimate 3-phosphate (EPSP) by competitively binding EPSP synthase (EPSPS) (38). In addition to plants, the shikimate pathway is also used by bacteria and some fungi. Bacteria with different EPSPS alleles encoding specific amino acid changes vary in their resistance to glyphosate, and can be classified as glyphosate resistant or sensitive based on their EPSPS allele (39). Bacteria can also be resistant to GBH through other mechanisms, such as exporting glyphosate out of the cell with efflux pumps (40) or by degrading it (41).

Here, we investigated how aquatic bacteria evolve in a semi-natural community faced with the GBH Roundup as an experimentally applied selective pressure. We exposed replicate mesocosms to two pulses of Roundup and sequenced metagenomes at five time points over eight weeks. To study evolutionary responses to selection imposed by GBH, we focused on 11 bacterial species (metagenome-assembled genomes; MAGs) present in multiple ponds after four weeks and tracked their genetic diversity in control and GBH-treated ponds. While we found some evidence for GBH- driven genome-wide selective sweeps in progress, these remained incomplete on the short timescale of our study and tended not to be repeatable across replicate ponds. We hypothesized that species predicted to be GBH-sensitive based on their EPSPS allele would be under stronger selection than GBH-resistant species. Although predicted sensitive and resistant species had no evident differences in genome-wide diversity after GBH treatment, they have distinct genetic targets of selection. Particularly in GBH-sensitive species, GBH selected for single nucleotide variants in genes involved in amino acid transport and metabolism, and for reduced copy number of genes involved in transcription and translation. Together, our results show how a known environmental stressor affects community diversity without consistently changing genome-wide within-species diversity beyond a few specific targets of selection.

## Results

### Quantifying within-species diversity across experimental treatments

In this experiment we sampled nine 1000 L ponds filled with pristine lake water over eight weeks to study how natural populations of aquatic bacteria evolve after exposure to Roundup, a commonly used GBH. Treated ponds received two pulses of GBH (at 15 mg/L and 40 mg/L) whereas controls received only phosphorus (as a control for the high phosphorus content of glyphosate) or no treatment. For each of the nine ponds, we co-assembled metagenomic reads from five time points throughout the 8-week experiment and binned the resulting contigs into metagenome-assembled genomes (MAGs), yielding a database of 315 non-redundant MAGs, which we define as distinct species (Methods). To quantify genetic diversity within species, we competitively mapped reads from each time point to our non-redundant MAG database and quantified the number and frequency of single nucleotide variants (SNVs) in each MAG in each experimental pond. We identified 11 MAGs that were present at a minimum of 4x average depth of coverage in at least one control and one GBH pond in the two samples collected at one and four weeks after the 15 mg/L GBH pulse (**Figure 1**). These 11 MAGs had an average size of 3.04 Mbp, completeness of 87.19%, and redundancy of 2.31% (**Table 1**). For some MAGs, SNV detection rates were correlated with MAG coverage (**Figure S1**). To resolve this bias, we subsampled reads such that each MAG had an equal coverage across ponds (Methods). The vast majority of these MAGs (10/11) were at low or undetectable relative abundance at time point 1, then increased at time point 2 in both control and treatment ponds (**Figure S2**). This suggests that this subset of species are well-adapted to the pond environment.

**Figure 1.**
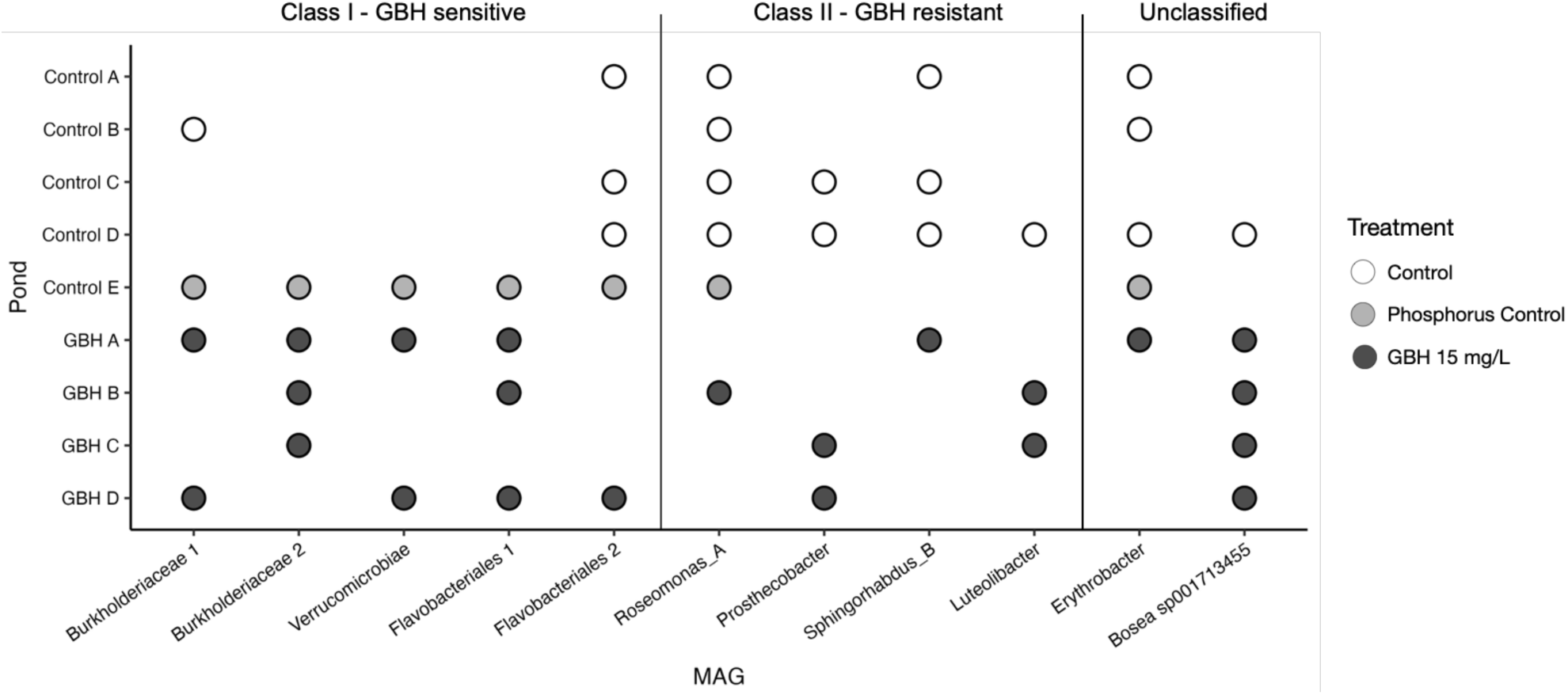
MAGs present in GBH treatment and control ponds after 15 mg/L GBH pulse. MAGs with a minimum of 4x average sequencing depth in both control and GBH ponds at time point 2 (merged samples from day 7 and 28). Ponds are coloured by the treatment received. MAGs are grouped by their predicted sensitivity to GBH based on their EPSPS allele.

**Table 1.**
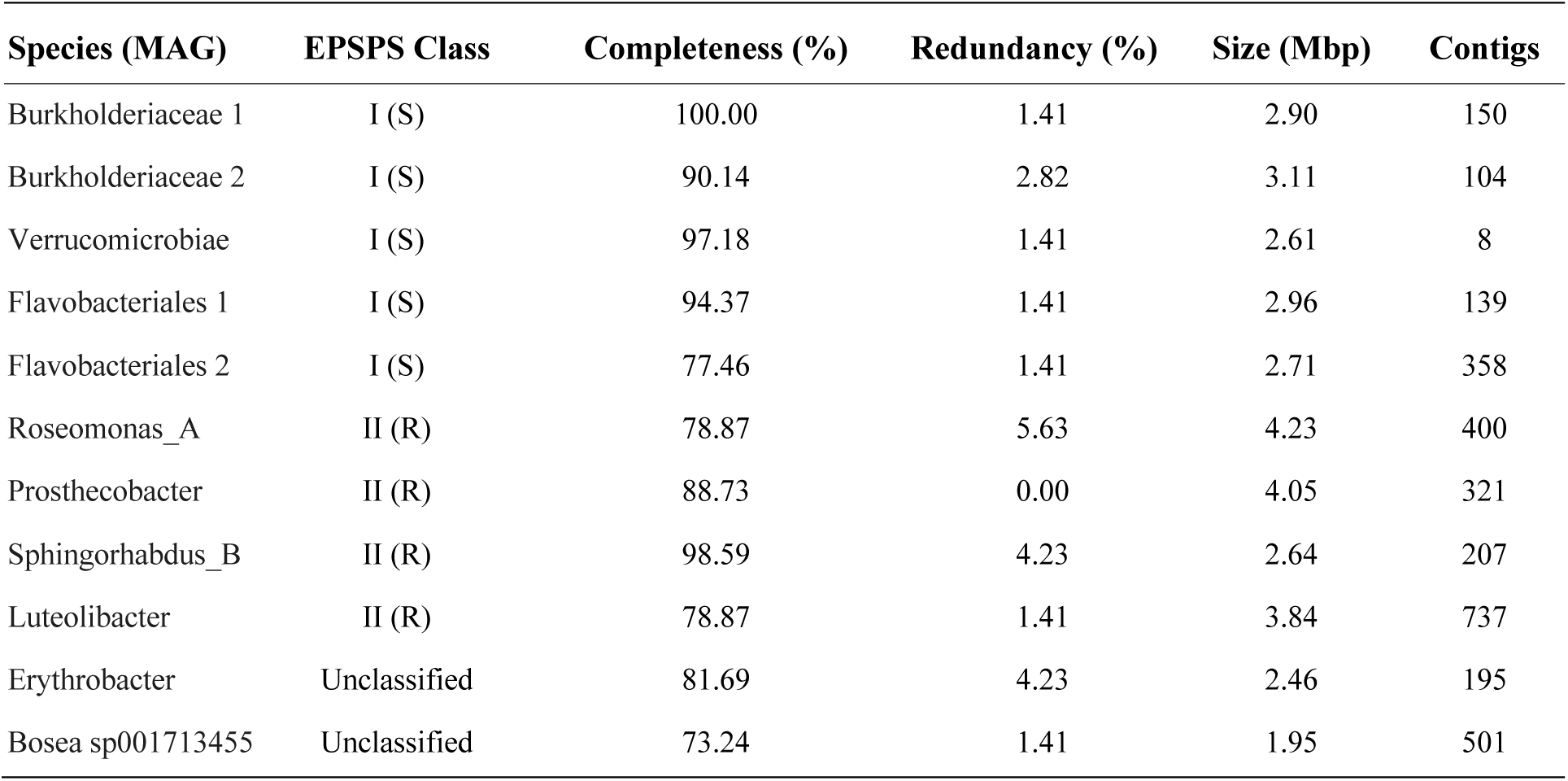
Summary of MAGs analyzed. Predicted sensitivity and resistance to glyphosate are indicated by (S) and (R) respectively.

If GBH inhibits the growth or viability of some bacterial species, we would expect GBH-treated ponds to have fewer detectable species (MAGs) present compared to control ponds. We estimated the species richness of each pond at time point 2 (after the 15 mg/L GBH pulse) by determining the number of MAGs in our database that were present. Consistent with expectation and with an earlier study (2), we found that GBH ponds had significantly lower species richness, approximately two-fold lower than control ponds (Wilcoxon Rank Sum test, U = 0, *p* = 0.016) (**Figure 2A**). The phosphorus control pond had lower richness than other controls, but still higher than any GBH- treated pond (**Figure 2A**). For most species, there was no evident relationship between within-species diversity and community diversity, with the exception of one *Verrucomicrobiae* MAG which showed a positive “diversity begets diversity” relationship (**Figure 2B**). This suggests that, on the time scale of our experiment, GBH has a significant effect on community-level diversity, but this effect is generally independent of within-species diversity.

**Figure 2.**
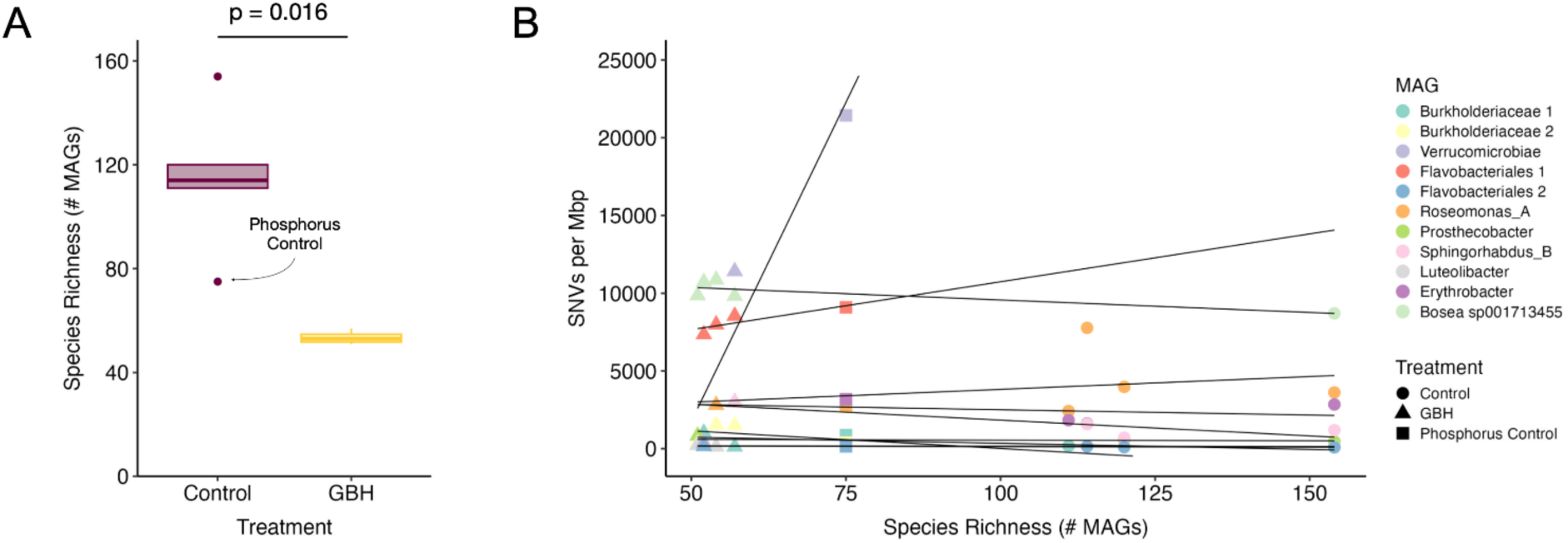
Species richness differs between GBH and control ponds independently of within-species diversity. The number of species (MAGs) present in each pond at time point 2. A species was considered present in a pond if the mean coverage was at least 1x and the breadth was > 50%. (A) Box plot of the number of species (MAGs) detected in each pond. (B) The relationship between within-species diversity (SNVs/Mbp) and species richness. A linear trendline was plotted for each MAG.

### Genome-wide selective sweeps are rare, incomplete, and not associated with EPSPS class

To study within-species diversity changes in greater detail, we compared SNV frequencies within each MAG between GBH and control ponds. If GBH drives genome-wide selective sweeps, this should result in genome-wide purges of genetic diversity in GBH-treated ponds compared to control ponds. One MAG, S*phingorhabdus_B*, classified as GBH-resistant, was the only species recovered in ponds both before and after the GBH pulse, allowing us to track genetic diversity over time. At time point 1, before any pond received GBH, the diversity in the *Sphingorhabdus_B* population varied somewhat between the five ponds (402 to 1198 SNVs/Mbp) (**Figure 3A & 3B, Table S1**). At time point 2, the diversity in control ponds increased slightly, but much more dramatically in GBH-treated pond A, reaching nearly 3,000 SNVs/MBp (**Figure 3A & 3B**). This increased diversity over time is not expected under a genome-wide selective sweep. Alternatively, such a pattern could be explained by a soft selective sweep in which beneficial mutations rise to fixation in multiple different genome backgrounds – for example, selecting for a rare strain with an adaptive allele to increase in frequency. Consistent with a soft sweep, this GBH pond had over 5000 fixed (reference allele frequency = 0) or nearly fixed substitutions (yellow in **Figure 3A**, **Table S1**) alongside the nearly 3000 polymorphic sites. In the other GBH replicate, S*phingorhabdus_B* was not recovered at time point 2, suggesting strong selection drove this species to extinction or below the limit of detection. *Sphingorhabdus_B* in the GBH A pond experienced approximately 4x more gene copy number changes over time than control ponds (**Figure S3, Table S2**). Further, 91% of genes with a copy number change in GBH A from T1 to T2 were also identified as having a copy number change between GBH A and control ponds at T2, compared to an average of 35% overlap among controls. This suggests that GBH is selecting for gene copy number changes in *Sphingorhabdus_B* over time. Together, these results suggest that natural selection imposed by GBH drives changes in genetic diversity over time, consistent with a soft or gene-specific sweep, or an incomplete genome-wide sweep that might have gone to completion given more time.

**Figure 3.**
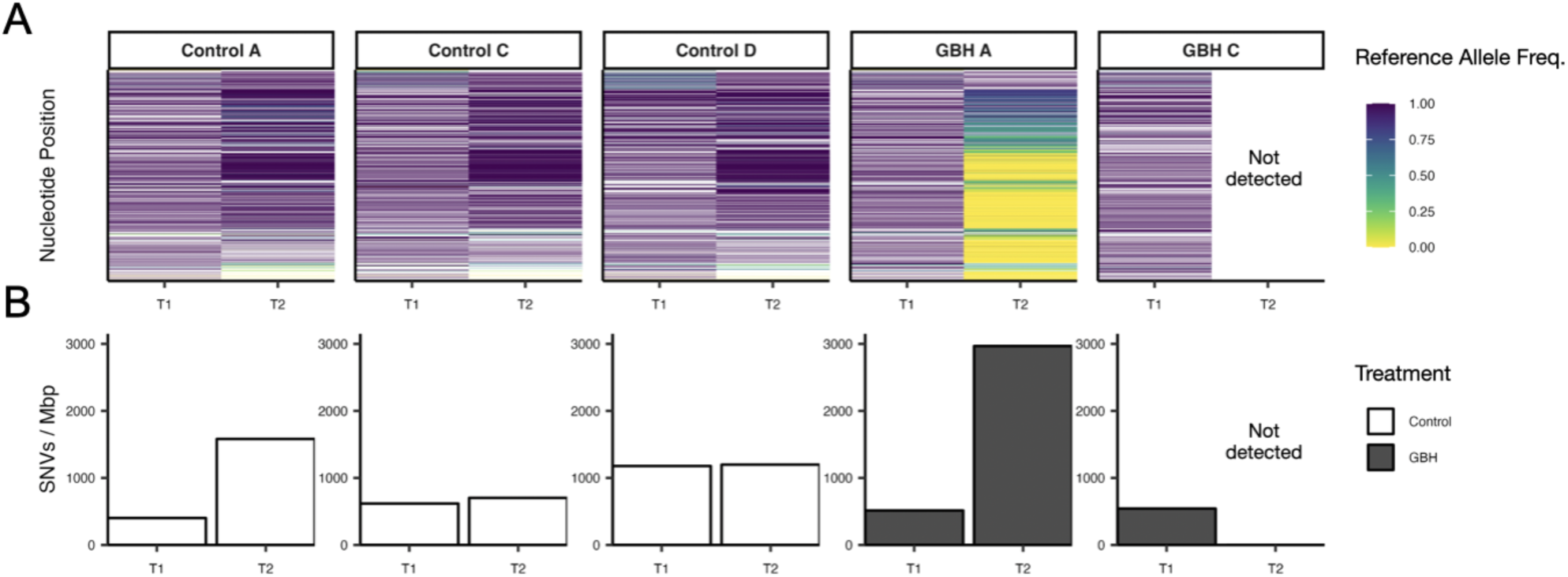
Temporal diversity changes in S*phingorhabdus_B* before and after a GBH pulse. (A) Each row in the heat map represents a nucleotide position which is polymorphic or fixed on a non-reference allele in at least one pond. The rows are coloured based on the reference allele frequency. Reference allele frequency at each genome position is calculated as the proportion of reads mapping to that site in the reference MAG that match the reference allele. Rows are ordered by the mean reference allele frequency across all ponds. (B) Total number of polymorphic sites in the MAG population divided by the MAG genome size (bp) x 10^6^. T1 (time point 1) includes the sample taken before any GBH pulse was added on day 0. T2 (time point 2) includes two samples taken after a 15 mg/L GBH pulse on days 7 and 28.

Although we lacked time series data for the other ten MAGs (which were too rare at T1 to call SNVs), we were still able to compare within-species diversity in control vs. GBH ponds at T2. Of these MAGs, none showed unequivocal evidence for a complete and repeatable GBH-driven genome-wide selective sweep (**Figure 4, Figure S3**). While in some cases there were differences in diversity between ponds, all populations contained measurable diversity in both control and GBH treatment ponds. In one example of a potential genome-wide selective sweep, the *Verrucomicrobiae* population had lower diversity in both GBH ponds compared to a control (**Figure 4A & 4B**). The *Verrucomicrobiae* population in the control pond contained 21,438 SNVs/Mbp, which was reduced by almost half in one GBH pond (11,410 SNVs/Mbp) and by over 50-fold in another (373 SNVs/Mbp; **Figure 4B, Table S1**). This is consistent with a genome-wide selective sweep in progress, favouring the reference allele. In the *Burkholderiaceae* 1 population, diversity was reduced in one replicate GBH pond but not the other (**Figure 4B**), suggesting a genome-wide sweep that was not repeatable, or was more rapid in one replicate. Both GBH ponds had more fixed non-reference alleles or low reference allele frequency SNVs compared to control ponds, suggesting a replacement of one dominant strain by another (**Figure 4A**). Similarly, a potential genome-wide sweep in the *Prosthecobacter* population only reduced diversity in one GBH pond compared to controls, resulting in the fixation of non-reference alleles (**Figure 4C & 4D).** The other GBH pond shared some of these fixed substitutions, but remained more diverse than either control pond (**Figure 4C & 4D)**.

**Figure 4.**
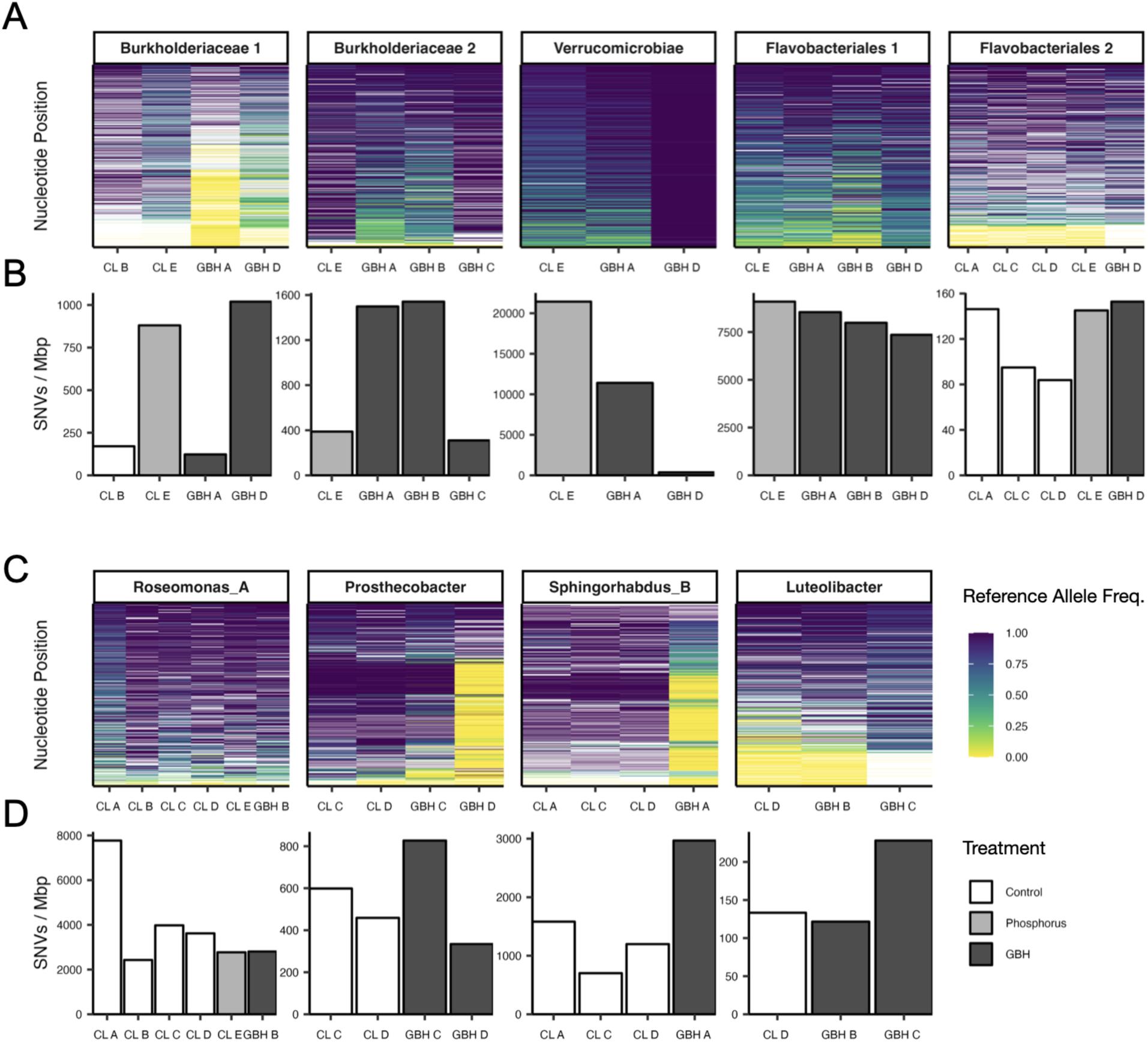
**Population diversity of GBH-sensitive and resistant MAGs**. (A & C) Each row in each heat map represents a nucleotide position which is polymorphic or fixed on a non-reference allele in at least one pond. The bars are coloured based on the reference allele frequency. Reference allele frequency at each genome position is calculated as the proportion of reads mapping to that site in the reference MAG that match the reference allele. Rows are ordered by the mean reference allele frequency across all ponds. (B & D) Total number of polymorphic sites in the MAG population divided by the MAG genome size (bp) x 10^6^. (A & B) Predicted GBH-sensitive MAGs. (C & D) Predicted GBH-resistant MAGs.

Overall, the evidence for genome-wide selective sweeps was equivocal. Defining a genome-wide selective sweep as a reduction in diversity across the genome in one GBH population compared to the average of the control populations, only 3/11 MAGs showed any evidence of a genome-wide selective sweep. We further hypothesized that GBH would impose stronger selection on MAGs predicted to be GBH-sensitive at the beginning of the experiment. We classified MAGs as GBH- sensitive or resistant based on their EPSPS allele. Of the 11 MAGs, five were classified as Class I (GBH-sensitive), four were classified as Class II (GBH-resistant), and two could not be classified because the EPSPS gene was not present in the annotation (**Figure 1**, **Table 1**). There were no apparent differences in genetic diversity after the GBH pulse between predicted GBH-sensitive MAGs (**Figure 4A & 4B**), resistant MAGs (**Figure 4C & 4D**), or unclassified MAGs (**Figure S3**), none of which showed evidence for complete, repeatable genome-wide selective sweeps driven by GBH.

### Identifying genetic targets of selection

Regardless of whether genome-wide selective sweeps occur, our data provide the opportunity to identify the likely genetic targets of GBH selection. It is typically challenging to identify the targets of selection after a genome-wide sweep, because the targets of selection are genetically linked to other hitchhiking mutations (42, 43). Our replicated study design alleviates this challenge because selected mutations could sweep independently in different replicate ponds (although this is not guaranteed to occur if the exact same genome fixes in replicate ponds). However, classic genome-wide sweeps appear to be rare in our experiment (**Figures 4 and S3**), and soft or gene-specific sweeps may be more common – both of which facilitate finding the targets of selection. We used four different approaches to identify genes targeted by GBH selection: (1) those with large changes in SNV frequency between GBH and control ponds, (2) those with consistently reduced SNV density between GBH and control ponds, (3) those with consistently reduced numbers of SNVs between GBH and control ponds, and (4) those with large changes in gene copy number between GBH and control ponds (Methods).

In the first approach, we calculated the difference between the mean reference allele frequency at each nucleotide position in control ponds and GBH ponds. We found that all populations had genes containing at least one large SNV frequency shift (range of 10 - 1,393 genes per MAG with a shift in reference allele frequency of 0.5 or more; **Table 2, Table S3**). Pooled across all MAGs, we found that these genes with shifts in SNV frequency were significantly enriched in several functional categories involved in metabolism (COG categories G, E, I, P, and Q; Fisher’s exact test, *p* < 0.05, FDR < 0.1; **Figure 5**). We also found a significant depletion of SNV frequency changes in genes from categories J (translation, ribosomal structure and biogenesis), K (transcription), N (cell motility), and X (mobilome). These results point to certain gene functions consistently targeted by selection across all populations.

**Figure 5.**
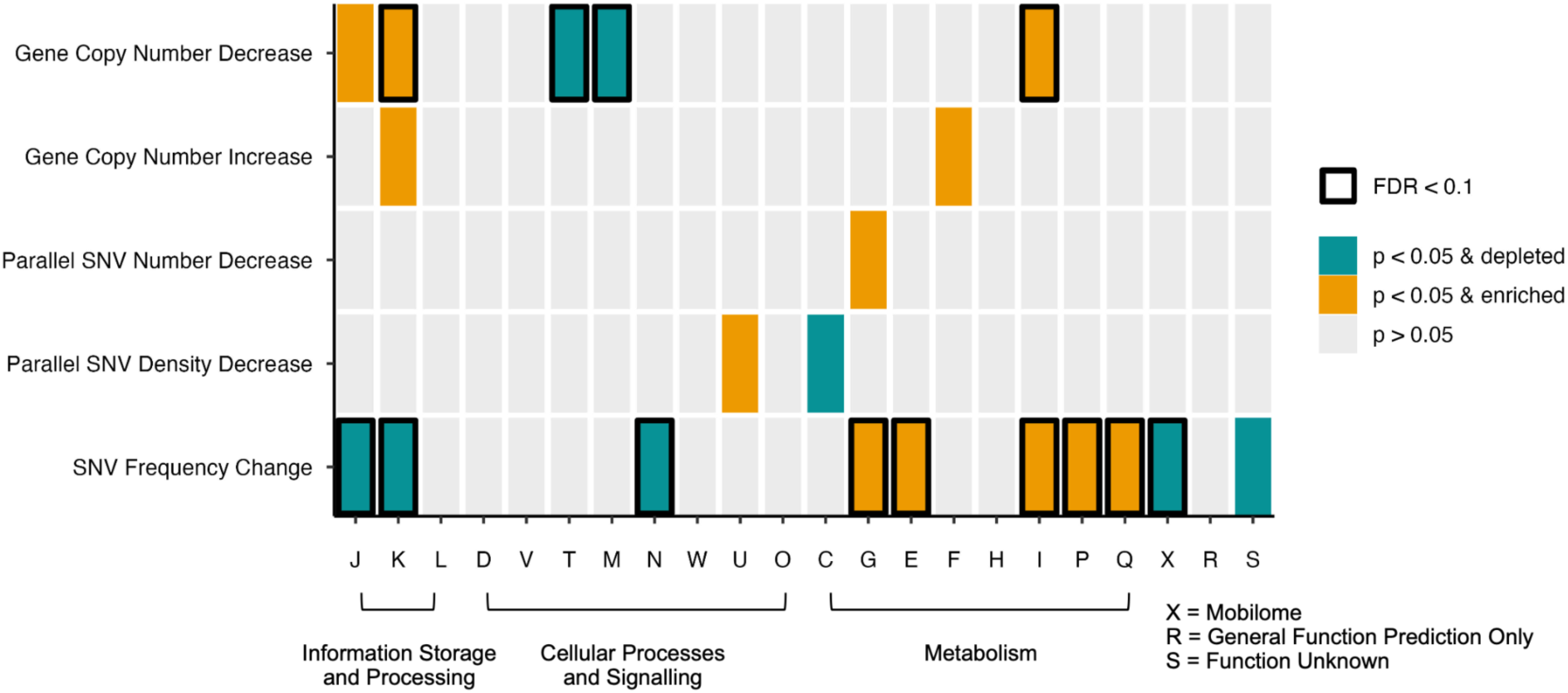
Functional categories enriched or depleted for genes identified as targets of selection. For each method used to identify possible targets of selection, a Fisher’s exact test was performed for each COG category between genes identified as targets of selection and all other genes present in the combined MAGs. Bars are coloured by significance and odds ratio ( < 1 = depleted or > 1 = enriched). Bars outlined in black pass our false discovery rate (FDR) threshold of < 0.1. COG categories: J = Translation, ribosomal structure and biogenesis, K = Transcription, L = Replication, recombination, and repair, D = Cell cycle control, cell division, chromosome partitioning, Y = Nuclear structure, V = Defense mechanisms, T = Signal transduction mechanisms, M = Cell wall/membrane/envelope biogenesis, N = Cell motility, Z = Cytoskeleton, W = Extracellular structures, U = Intracellular trafficking, secretion, and vesicular transport, O = Posttranslational modification, protein turnover, chaperones, C = Energy production and conversion, G = Carbohydrate transport and metabolism, E = Amino acid transport and metabolism, F = Nucleotide transport and metabolism, H = Coenzyme transport and metabolism, I = Lipid transport and metabolism, P = Inorganic ion transport and metabolism, Q = Secondary metabolites biosynthesis, transport and catabolism, X = Mobilome: prophages, transposons, R = General function prediction only, S = Function unknown Lastly, we identified genes that differed in copy number between control and GBH ponds. For each gene, we compared its relative depth of coverage in controls to GBH ponds (Methods). We found that all populations had genes that increased or decreased by > 0.5 gene copies between control and GBH ponds (**Table 2, Table S6**). Genes that decreased in copy number were significantly enriched in transcription (K) and lipid transport and metabolism (I) and depleted in signal transduction (T) and cell wall/membrane biogenesis (M) (Figure 5).

**Table 2.**
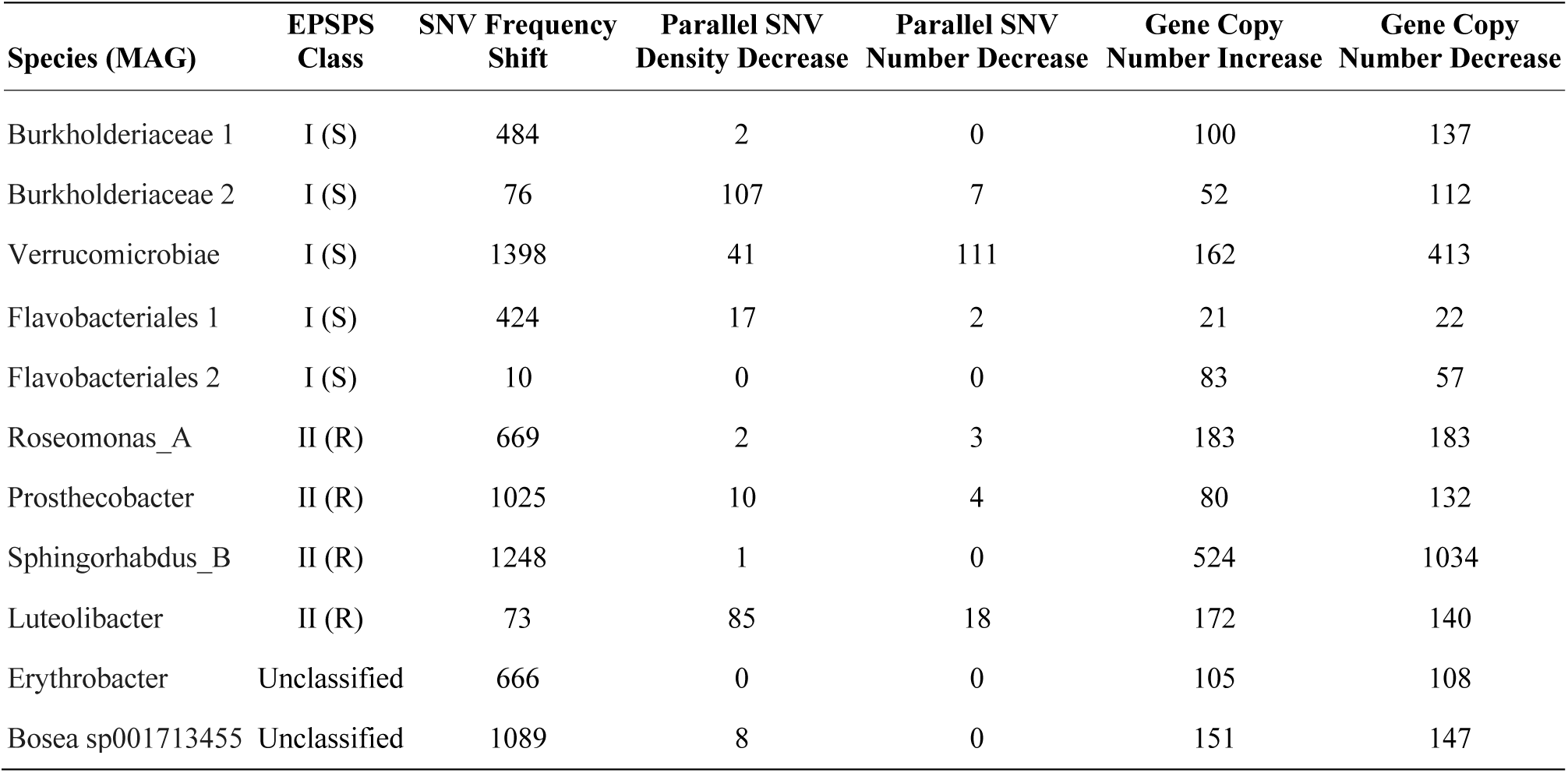
Summary of the number of genes identified as targets of selection. Predicted sensitivity and resistance to glyphosate are indicated by (S) and (R) respectively.

Next, we identified genes in GBH-exposed populations with less diversity compared to the rest of the genome, as expected in a gene targeted by a selective sweep. Nine of the 11 populations had at least one gene (range of 1 - 107) with a repeatable (parallel across replicate ponds) decrease in SNV density in each GBH-control comparison (**Table 2, Table S4**). Further, six of these populations had at least one gene (range of 2 - 111 genes) with a significant SNV number decrease in each GBH-control comparison using an existing method (44) which identifies genes with differential SNV counts between treatments (**Table 2, Table S5**). These parallel decreases in both SNV density and number provide strong evidence for selection since they occur repeatedly across replicates. These genes were enriched in carbohydrate transport and metabolism and intracellular trafficking, secretion, and vesicular transport (COG categories G and U, Fisher’s exact test, *p* < 0.05). Even if not significant after multiple hypothesis correction (FDR > 0.1), COG category G was also enriched in large SNV frequency changes, supporting its importance as a target of GBH selection (**Figure 5**).

Overall, genes involved in metabolism were frequently identified as targets of selection by different methods, while genes involved in cellular processes and signalling were rarely identified (**Figure 5**). Meanwhile, translation and transcription (J and K) tended to decrease in gene copy number under GBH treatment. Selection on amino acid transport and metabolism, along with reduced copy number of transcription and translation genes, is mainly driven by MAGs classified as GBH-sensitive (**Figure S4**). This could be due to GBH-sensitive species, but not resistant species, being under selection for changes in amino acid metabolism – which contains the shikimate pathway targeted by glyphosate. Thus, MAGs with inferred GBH resistance or sensitivity at the beginning of the experiment may have experienced different genetic targets of selection.

## Discussion

Previous work has shown GBHs like Roundup alter the composition of aquatic bacterial communities (2, 36). However, how individual bacterial populations within a community evolve in response to GBH is unknown. More generally, our understanding of microbial evolution in nature has been hindered by the lack of studies combining experimentally controlled selective pressures with measures of within-species diversity (evolution) in the context of a diverse community (45). To fill this gap, we quantified within-species diversity in bacterioplankton exposed to GBH in controlled setup freshwater mesocosm ponds. We found that genome-wide selective sweeps were rare and often incomplete over the time scale of the experiment. Rather, all of the 11 populations studied contained measurable and often substantial genetic diversity (on the order of hundreds to thousands of SNVs per Mbp, consistent with standing strain-level diversity) in both GBH-treated and control ponds. Potential selective sweeps in one pond were often not observed in a replicate pond. This lack of repeatability raises doubt about whether a sweep was driven by selection or drift.

There are several explanations for the lack of genome-wide selective sweeps documented in this study, which are not mutually exclusive. First, a four-week time scale might have been too short to observe sweeps proceeding to completion. One other potential genome-wide sweep documented in a lake took years to complete (31) so it is plausible that sweeps in natural environments require much longer to complete than the few dozen generations shown in simplified evolutionary models (46). Second, it is possible that the community-level response to selection is dominated by species sorting, with pre-existing GBH-resistant taxa flourishing and sensitive taxa declining, giving little opportunity for evolutionary responses within these species. A previous experiment in the same mesocosm system showed that GBH substantially restructures the bacterioplankton community and reduces diversity (2), which we confirmed in this study. It is therefore plausible that ecological responses dominated evolutionary responses in this context and time scale. Third, recombination-driven gene-specific selective sweeps or soft selective sweeps may be more common than genome-wide selective sweeps in our experiment. We detected signals of selection in numerous genes, including parallel purges of diversity in GBH-treated ponds but not controls. Gene-specific or soft selective sweeps are plausible mechanisms to explain these patterns. Using short-read metagenomic data, it is difficult to infer recombination; therefore direct evidence for recombination-driven gene-specific selective sweeps is lacking. Given the relatively high levels of within-species standing genetic diversity, which is typical in a natural community, adaptive mutations could spread on distinct genetic backgrounds (“strains”) in a soft selective sweep, resulting in the maintenance of diversity without high levels of recombination. Future work using long-read sequencing or whole genomes from isolated bacteria will be needed to resolve the relative influence of soft and gene-specific selective sweeps.

Despite the lack of evident genome-wide selective sweeps, we identified genes with consistent changes in allele frequency or copy number in GBH ponds, consistent with natural selection acting on specific cellular pathways in multiple species. Glyphosate binds EPSPS and prevents aromatic amino acid synthesis; we therefore expected natural selection on EPSPS and surrounding amino acid metabolic pathways. We classified each species as putatively resistant or sensitive to glyphosate at the beginning of the experiment based on known mutations in the EPSPS gene. We expected GBH-sensitive populations to be under stronger selection than GBH-resistant populations, but we did not observe any notable differences in their genome-wide diversity between treatments. However, we did find evidence that the functional categories of genes under selection differed between GBH-sensitive and resistant populations. Although *aroA*, the gene encoding EPSPS, was not identified as a target of selection in any species, genes involved in amino acid metabolism (COG category E) were significantly enriched in SNV frequency changes in response to GBH across all species – an effect that is driven by GBH-sensitive species. This suggests that amino acid metabolism pathways surrounding EPSPS might be specifically under selection in species with a GBH-sensitive EPSPS allele. Other GBH-sensitive species might be selected for slower growth, as has been observed in *B. subtilis* facing nutrient starvation (47). As evidence for this, we found that GBH selected for a decreased copy number of transcription and translation genes in GBH-sensitive MAGs, but not in GBH-resistant MAGs. Together with previous findings that GBH selects for multidrug efflux pumps in bacterioplankton communities (36), our results highlight that targets of selection can be diverse in natural communities, not always centered on canonical resistance mutations.

Why *Prosthecobacter* and *Sphingorhabdus_B* — both GBH-resistant MAGs based on their EPSPS allele — would be targeted by GBH selection is unclear. The *Prosthecobacter* population experienced fixation of many alternate alleles in one GBH replicate but not the other. Such an inconsistent response suggests the dominance of drift over GBH-driven selection. In *Sphingorhabdus_B* where the starting population at time point 1 is known, the number of SNVs (including many at low frequency) increased along with the number of fixations in one GBH replicate. One hypothesis is that GBH interferes with the cell in unknown ways, beyond the canonical EPSPS target. Alternatively, selection on GBH-resistant species could be indirect. For example, GBH causes shifts in community composition, which could subsequently impose new selective pressures on these species through competition, cross feeding, or by creating novel niches (8, 19). Importantly, the potential soft sweep in *Sphingorhabdus_B* was only observed in one pond, with the MAG decreasing below the limit of detection in the other pond after the GBH pulse. This again suggests that stochastic dynamics (drift) could dominate selection in a species close to the limit of local extinction.

Further research will be needed to quantify selective sweep dynamics over longer time scales, and for a larger number of taxa. Ten of the MAGs analyzed were present at very low or undetectable abundances at time point 1, preventing us from tracking changes in diversity over time. This is unlikely to affect our conclusions regarding genome-wide selective sweeps (or the lack thereof) because each pond was filled with water from the same lake at the same time, making it improbable that the initial diversity within each MAG varied substantially across ponds. Nevertheless, additional time-series – over longer time scales – would provide more robust support for our conclusions. With bacterial doubling in nature roughly every 1-25 hours (48), the pond populations likely evolved for approximately 28-700 generations after the GBH pulse. Depending on the strength of selection, this could be sufficient time to detect changes in allele frequencies (including fixation) for fast-growing, but perhaps not slow-growing species. While the 11 populations we tracked encompass both GBH-sensitive and resistant EPSPS classes, they are far from representing the full bacterial diversity present in Lac Hertel. Unfortunately, many MAGs that were recovered in multiple control ponds and time points were absent from GBH ponds after the GBH pulse, which prevented us from including them in the analysis. This observation is likely due to the selective pressure imposed by GBH, where MAGs sensitive to GBH drastically decrease in abundance after the GBH pulse and are not captured at our sequencing depth. While we do not see strong evidence for complete genome-wide selective sweeps across the 11 populations we tracked, it remains unknown if sweeps occurred in any of the other 304 populations that were too rare to detect using our shotgun metagenomic approach.

Our study represents an advance in our understanding of evolution in natural environments. By applying a known selective pressure to semi-natural communities, we were able to go beyond documenting sweep-like patterns with unknown causes, and move toward attributing these patterns to selection. In our limited sample of species, and over short time scales, we can conclude that genome-wide selective sweeps are rare in response to GBH, a stressor with clear community-level effects. Even if genome-wide sweeps are rare, we show that GBH imposes selection on several categories of genes, including those involved in amino acid metabolism, transcription and translation – particularly in GBH-sensitive species. This highlights the potentially unexpected evolutionary consequences of agrochemical runoff into freshwater ecosystems. Future work will be needed to determine if genome-wide selective sweeps are more common in response to different selective pressures, in rarer taxa, and over longer time scales.

## Methods

### Experimental design, sampling, and sequencing

In this experiment, nine “ponds” (mesocosms) were filled with approximately 1000L of water originating from Lac Hertel, a pristine lake with no known prior herbicide or pesticide contamination, at Gault Nature Reserve in Mont-Saint-Hilaire, Quebec. We sampled the nine ponds five times over 8-weeks between July 16th, 2021 and September 10th, 2021. Four out of nine ponds received two pulses of the glyphosate-based herbicide, Roundup, in the form of Roundup Super Concentrate Grass and Weed Control (reg. no. 22759; Bayer). One control pond received a single pulse of phosphorus (K2PO4) to control for Roundup as a phosphorus source (49). The four remaining control ponds did not receive any treatment. On day 0, the four GBH ponds received a 15 mg/L GBH pulse and the phosphorus control pond received a 320 μg/L pulse of phosphorus (K2PO4). On day 28, the four GBH ponds received a 40 mg/L GBH pulse. GBH concentrations were calculated based on the concentration of glyphosate acid. We collected 1L of water at five timepoints: day 0 (pre-GBH pulse), 7 & 28 (after 15 mg/L GBH pulse), and 35 & 56 (after 40 mg/L GBH pulse). For each sample, we filtered 250 mL of water through a 0.22µm pore size polyethersulfone membrane (Sigma-Aldrich) to collect the bacterial community for metagenomic sequencing. The filters were stored at -80 **°**C prior to DNA extraction. DNA was extracted using the DNeasy PowerWater Kit (QIAGEN), libraries were prepared with the NEBNext Ultra II DNA Library Prep kit (New England Biolabs), and sequenced on an Illumina NovaSeq6000 S4 v.1.5 with 150bp paired-end reads.

### Metagenomic assembly, binning, and classification of metagenome-assembled genomes (MAGs)

Metagenomic reads were trimmed with trimmomatic v0.39 (50) to remove illumina adapters and discard low quality reads. Next, we co-assembled metagenomic reads by pond, pooling the five timepoints for each pond, with MEGAHIT v1.2.9 (51, 52). Following the anvi’o metagenomics workflow, we used anvi’o v7 (53) to generate a contig database for each pond’s co-assembly, identify genes with Prodigal v2.6.3 (54), and annotate each database with single copy gene taxonomy with HMMER (55). Next, metagenomic reads from each timepoint were mapped to the co-assembly of their corresponding pond with bowtie2 v2.4.4 (56) using default parameters. Samtools v1.15 (57) was used to convert the SAM output to BAM format and sort and index the BAM file. We used anvi’o to profile contigs longer than 2500 bp in each pond’s contig database by calculating coverage and single nucleotide variants across samples using the BAM files. We created a merged contig profile for each pond and used CONCOCT v1.0.0 (58) to bin contigs based on nucleotide composition, kmer frequencies, and coverage across samples. This resulted in 1536 bins across the nine ponds. Bins with > 70% completion were manually refined using the anvi’o interactive interface to remove contigs that were incorrectly binned (53). The 606 bins that had > 70% completion and < 10% redundancy were considered MAGs and were dereplicated at a 98% average nucleotide identity (ANI) with anvi’o’s dereplication command using fastANI v1.32 (59) resulting in 315 non-redundant MAGs. Taxonomic classification of MAGs was done using the Genome Taxonomy Database (GTDB) Release 214 (60) with GTDB-Tk v2.1.0 (61, 62).

### Gene prediction, annotation, and SNV calling

We used prodigal v2.6.3 (54) in metagenomic mode to predict protein coding genes in our database of 315 non-redundant MAGs. Functional annotation of genes was done using eggNOG mapper v2.1.12 (63, 64) using DIAMOND (65) to align sequences. MAGs with an annotated EPSPS gene (*aroA*) were classified as class I (GBH-sensitive) or class II (GBH-resistant) using the online classifier “*EPSPSClass* server” (39) which looks for specific amino acid markers within the gene. Next, metagenomic reads from each timepoint and pond were competitively mapped to our MAG database using bowtie2 v2.4.4 (56) with default parameters. Mapping outputs were converted to BAM format and indexed with samtools v1.16.1 (57). To increase the coverage for SNV calling, BAM files from the two time points after the 15 mg/L GBH pulse (day 7 and day 28) were merged using samtools v1.16.1 (57) and are henceforth referred to as time point 2. Similarly, the two time points after the 40 mg/L pulse (day 35 and day 56) were merged and are referred to as time point 3. For each MAG used for further analysis, to control for coverage bias in SNV detection, we subsampled mapped reads with samtools v1.18 (57) to match the coverage of the lowest coverage pond. We used inStrain v1.8.0 (66) to identify SNVs in each MAG population at the three time points. We set a minimum MAPQ of 2 to discard multi-mapped reads, a minimum ANI of reads mapping to the reference database of 95%, and a minimum position coverage of 5x to call a SNV. We discarded SNVs within 100 bp of contig edges and SNVs with a position depth greater than 3 times the MAG coverage or less than ⅓ of the MAG coverage. These depth filters remove both low-coverage regions and high-coverage regions containing likely mismapped reads, using established cutoffs (8, 67). We further removed SNVs with more than two alternative alleles. The inStrain command “IS.get(‘covT’)” was used to extract the depth at each position to differentiate between positions where the reference frequency was 1 (and thus depth at that position was not reported in the output) and positions with a depth too low to call a SNV (< 5x).

### Species richness

To estimate the species richness in each pond after the 15 mg/L GBH pulse, we first subsampled metagenomic reads from each pond with samtools v1.18 (57) to match the pond with the lowest number of reads at time point 2. Next, we used CoverM (68) to estimate the number of MAGs in our database of 315 MAGs that were present with at least 1X average depth and a 50% breadth of coverage. We performed a Wilcoxon Rank Sum test in R version 4.3.3 (69) to determine if GBH treated ponds had lower species richness (number of MAGs present) than control ponds. For each MAG with at least 4x coverage, we plotted the number of polymorphic sites in that MAG versus the species richness for each pond the MAG was present in. To determine if higher species richness in a pond is correlated with higher polymorphic sites within a MAG, we plotted a linear trendline for each MAG.

### Identifying genes with differential shifts in SNV frequency between treatments

We identified genes with large shifts in SNV frequency between GBH and control ponds. For each SNV, we calculated the mean reference frequency in control ponds and the mean reference frequency in GBH ponds. We calculated the difference between reference allele frequencies in control and GBH ponds, and considered as ‘large’ any absolute difference greater or equal to an arbitrary threshold of 0.5. These SNVs were further filtered to remove SNVs where there was overlap in the reference allele frequency between control and GBH ponds: if the average GBH pond reference allele frequency is low compared to controls, all GBH ponds must have a reference allele frequency lower than all control ponds. This assures that the direction of allele frequency change is consistent across treatments. Next, we determined which genes contain at least one SNV with a differential shift of at least 0.5 in SNV frequency between treatments. Note that some of these genes may contain SNVs with opposing directional shifts.

### Identifying genes with a decrease in diversity between treatments

Genes experiencing sweeps are expected to be purged of genetic diversity as an adaptive allele rises in frequency. To identify such genes, we focused on those with a reduced number of SNVs during GBH treatment compared to controls. We identified genes with fewer SNVs regardless of their frequency. For this analysis, we only included genes with at least 3x coverage in every pond and excluded genes with a coverage greater than 3 times the MAG coverage as this could be from read donating from other species which share the same gene (67). We calculated the SNVs density (per Mbp) for each gene and then calculated the difference in each pond-pond comparison. For each comparison, we selected the top 5% of genes which have a decrease in SNV density. From this list we selected genes which have a SNV density decrease in each GBH-control comparison (which we term a parallel SNV density decrease). We removed genes that were in the top 5% of any GBH-GBH or control-control comparison since these should not be relevant to GBH adaptation.

As a complement to this analysis, we used an existing method which identifies genes with differential SNV counts between treatments (44). This method assumes a Poisson distribution of mutations among genes (44). Briefly, for each MAG, we compared the total number of SNVs in a gene in one pond to the number of SNVs in that gene in another pond. For each pair of ponds, we identified genes with a significant SNV number difference between ponds. Next, we determined which genes had a significant decrease in SNV number in GBH ponds compared to controls in every GBH-control comparison (which we term a parallel SNV number decrease).

### Gene copy number variation

For each gene, we estimated the number of gene copies to identify gene copy number differences between treatments. Gene copy number was calculated as the gene’s depth of coverage divided by the average MAG depth of coverage. Genes with a copy number greater than 3 (likely enriched in mismapped reads) in any pond were excluded from the analysis. For each remaining gene we compared the average copy number in control ponds to the average copy number in GBH ponds. We considered genes with a copy number difference of at least 0.5 between control and GBH ponds to be notable. We further removed genes if there was any overlap in copy number between any control and GBH pond (e.g. if GBH copy number average is lower than the control average, all GBH ponds had to have a copy number lower than all control ponds).

### COG function enrichment analysis

We performed a gene set enrichment analysis using the COG database (70) on the five sets of genes identified as potential targets of selection: (1) SNV frequency changes, (2) parallel SNV density decreases, (3) parallel SNV number decreases, (4) gene copy number increases, and (5) gene copy number decreases. Genes in each MAG were annotated with COG IDs using eggNOG mapper v2.1.12 (63, 64) and COG categories from the 2020 database update (71). We used R version 4.3.3 (69) to perform a Fisher’s exact test with each gene list to determine if any COG category was significantly enriched for potential targets of selection compared to a background set of all genes from the 11 MAGs. A false discovery rate (FDR) correction was applied to account for multiple comparisons. In addition to the overall enrichment test, we performed separate tests for inferred GBH-sensitive and resistant MAGs.

## Data Availability

Metagenomic sequences from each sample are available on NCBI under BioProject PRJNA1161687.

## Code Availability

Scripts for all analyses are available at https://github.com/emderrick/LEAP_sweeps.

## Acknowledgements

We thank Delaney Barth for her assistance in the field and in the lab, Michelle Gros for her assistance in the field, the McGill Genome Centre sequencing platform, and members of the Shapiro lab for their feedback on the project. This study was supported by an NSERC Discovery Grant to B.J.S. LEAP was constructed under the leadership of Andrew Gonzalez, with funding from the Canadian Foundation for Innovation for which we are grateful.

## Notes

### Competing Interest Statement

The authors have declared no competing interest.

