## Supplementary Figures for "A glyphosate-based herbicide selects for genetic changes while retaining within-species diversity in a freshwater bacterioplankton community"

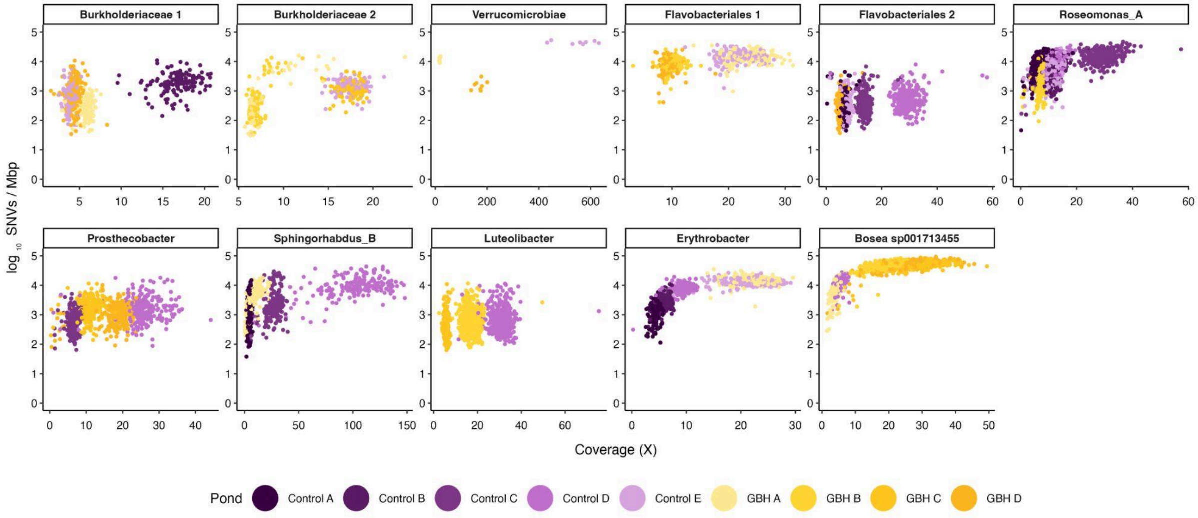


**Supplementary Figure S1. SNV detection correlates with MAG coverage for some MAGs.** Each dot in the graph represents one MAG contig. SNVs/Mbp is calculated as the number of polymorphic sites in the contig divided by the contig size * 10^6.

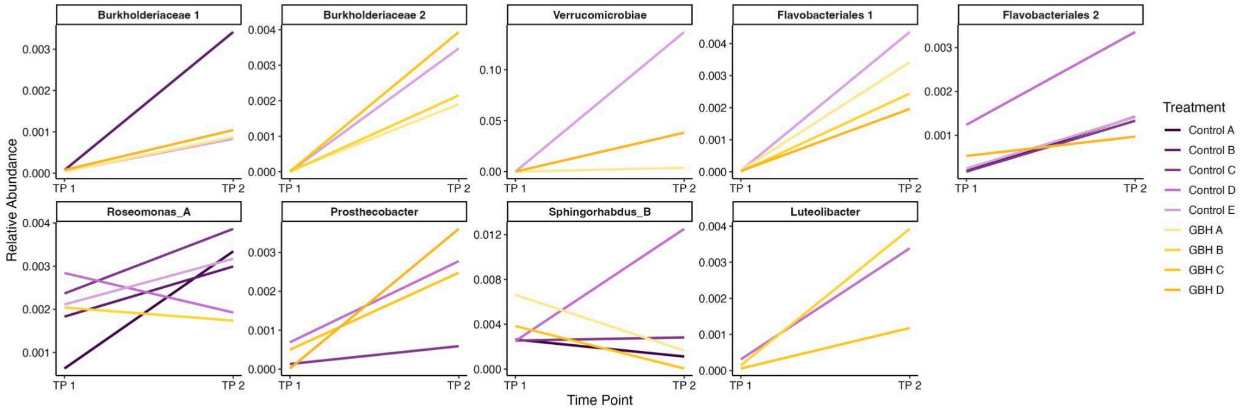


**Supplementary Figure S2. Relative abundance of MAGs before and after a 15 mg/L GBH pulse.** Relative Abundance is calculated as the number of reads mapping to a MAG / the total number of reads in sample / genome length (bp) * 10^6.

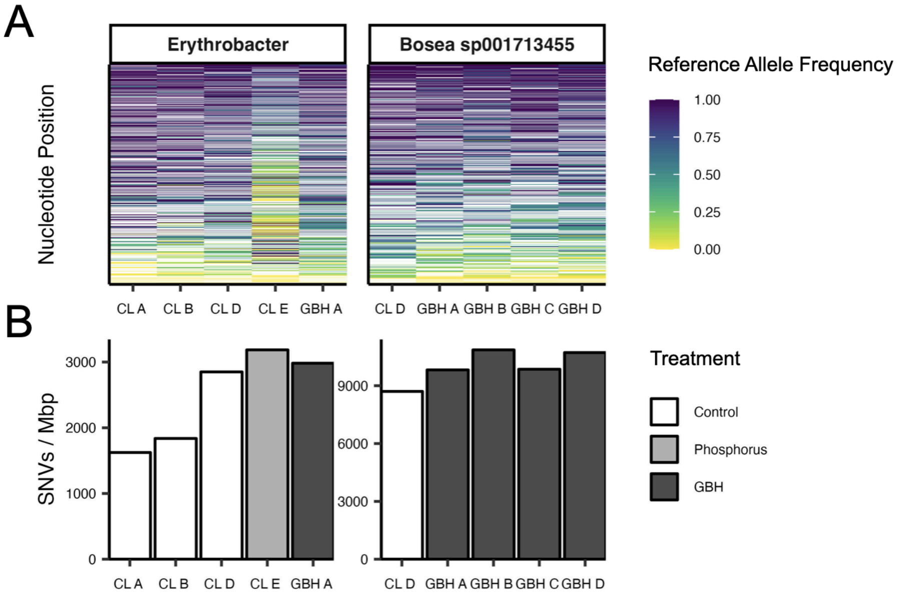


**Supplementary Figure S3. Population diversity of unclassified GBH sensitivity MAGs.** (A) Each row in each heat map represents a nucleotide position which is polymorphic or fixed on a non-reference allele in at least one pond. The bars are coloured based on the reference allele frequency. Reference allele frequency at each genome position is calculated as the proportion of reads mapping to that site in the reference MAG that match the reference allele. Rows are ordered by the mean reference allele frequency across all ponds. (B) Total number of polymorphic sites in the MAG population divided by the MAG genome size (bp) x $10^6$.

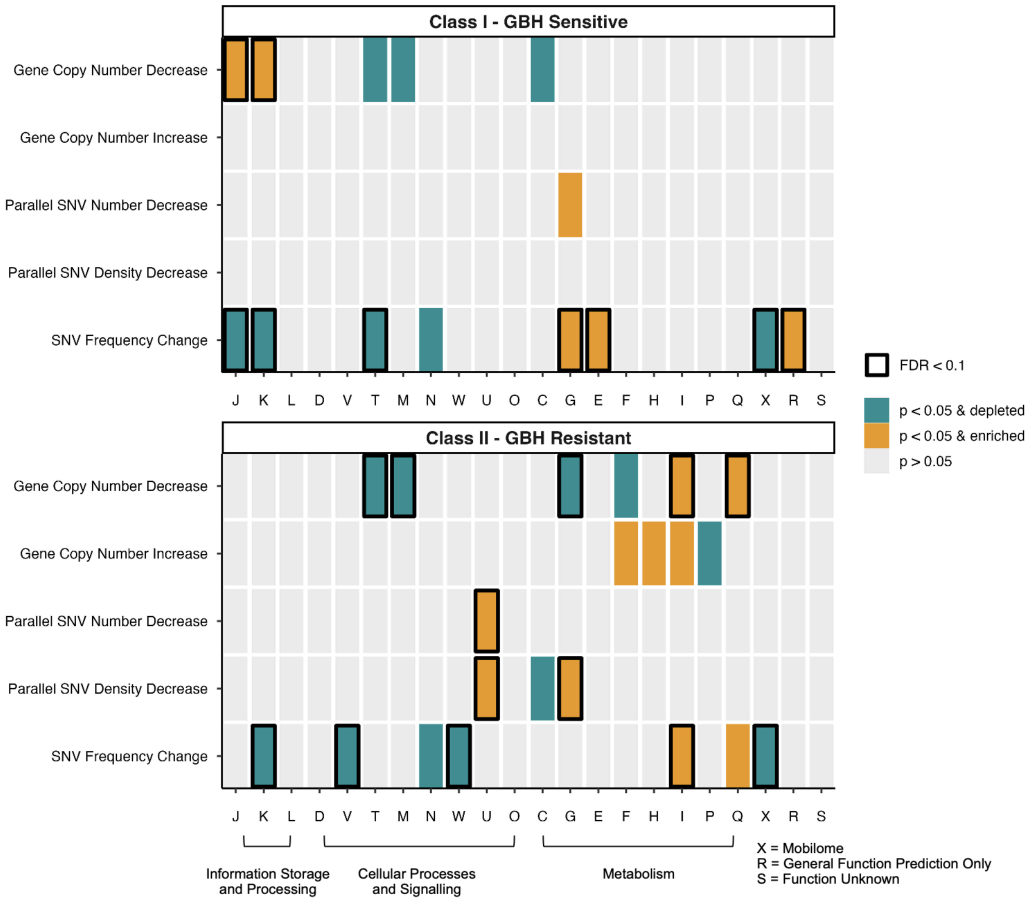


**Supplementary Figure S4. Functional categories enriched or depleted for genes identified as targets of selection separated by MAG predicted GBH sensitivity.** For each method used to identify possible targets of selection, a Fisher's exact test was performed for each COG category between genes identified as targets of selection and all other genes present in the combined MAGs. Bars are coloured by significance and odds ratio ( < 1 = depleted or > 1 = enriched). Bars outlined in black pass our false discovery rate (FDR) threshold of < 0.1. COG categories: J = Translation, ribosomal structure and biogenesis, K = Transcription, L = Replication, recombination, and repair, D = Cell cycle control, cell division, chromosome partitioning, Y = Nuclear structure, V = Defense mechanisms, T = Signal transduction mechanisms, M = Cell wall/membrane/envelope biogenesis, N = Cell motility, Z = Cytoskeleton, W = Extracellular structures, U = Intracellular trafficking, secretion, and vesicular transport, O = Posttranslational modification, protein turnover, chaperones, C = Energy production and conversion, G = Carbohydrate transport and metabolism, E = Amino acid transport and metabolism, F = Nucleotide transport and metabolism, H = Coenzyme transport and metabolism, I = Lipid transport and metabolism, P = Inorganic ion transport and metabolism, Q = Secondary metabolites biosynthesis, transport and catabolism, X = Mobilome: prophages, transposons, R = General function prediction only, S = Function unknown
